# Protocol for studying membrane protein dynamics and associated synaptic vesicle recruitment on native membrane sheets

**DOI:** 10.64898/2026.07.02.736009

**Authors:** Akshay Kapadia, Anne-Sophie Hafner

**Affiliations:** Donders Institute for Brain, Cognition and Behavior, Radboud University; Nijmegen, Netherlands; Interdisciplinary Institute for Neuroscience (IINS), Université de Bordeaux, UMR 5297 CNRS, 33076 Bordeaux, France; Technical contact

## Abstract

Plasma membrane sheets generated by controlled mechanical disruption provide direct access to the cytosolic face of the plasma membrane while preserving the native organization of membrane-associated proteins and lipids. Here, we present a protocol for generating and validating sonication-derived plasma membrane sheets from cultured cells, primary neurons, and isolated synaptosomes. We further describe their application for live and fixed imaging of membrane protein localization, organization, conformational dynamics, and protein-protein interactions, as well as quantitative membrane-associated synaptic vesicle recruitment assays.

This versatile platform preserves the native membrane environment while enabling direct visualization and quantitative analysis of membrane-associated processes at high spatial resolution. The protocol can be readily adapted to investigate diverse membrane proteins, lipid-dependent mechanisms, and vesicle tethering events across a wide range of cellular systems.

**Highlights:**

- Preparation of plasma membrane sheets exposing the cytosolic membrane surface
- Optimization and validation of membrane sheets from cultured cells, neurons, and isolated synaptosomes
- Adaptable platform for studying membrane protein interactions in native membranes
- Quantitative assay for membrane-associated synaptic vesicle recruitment

**Graphical Abstract:** 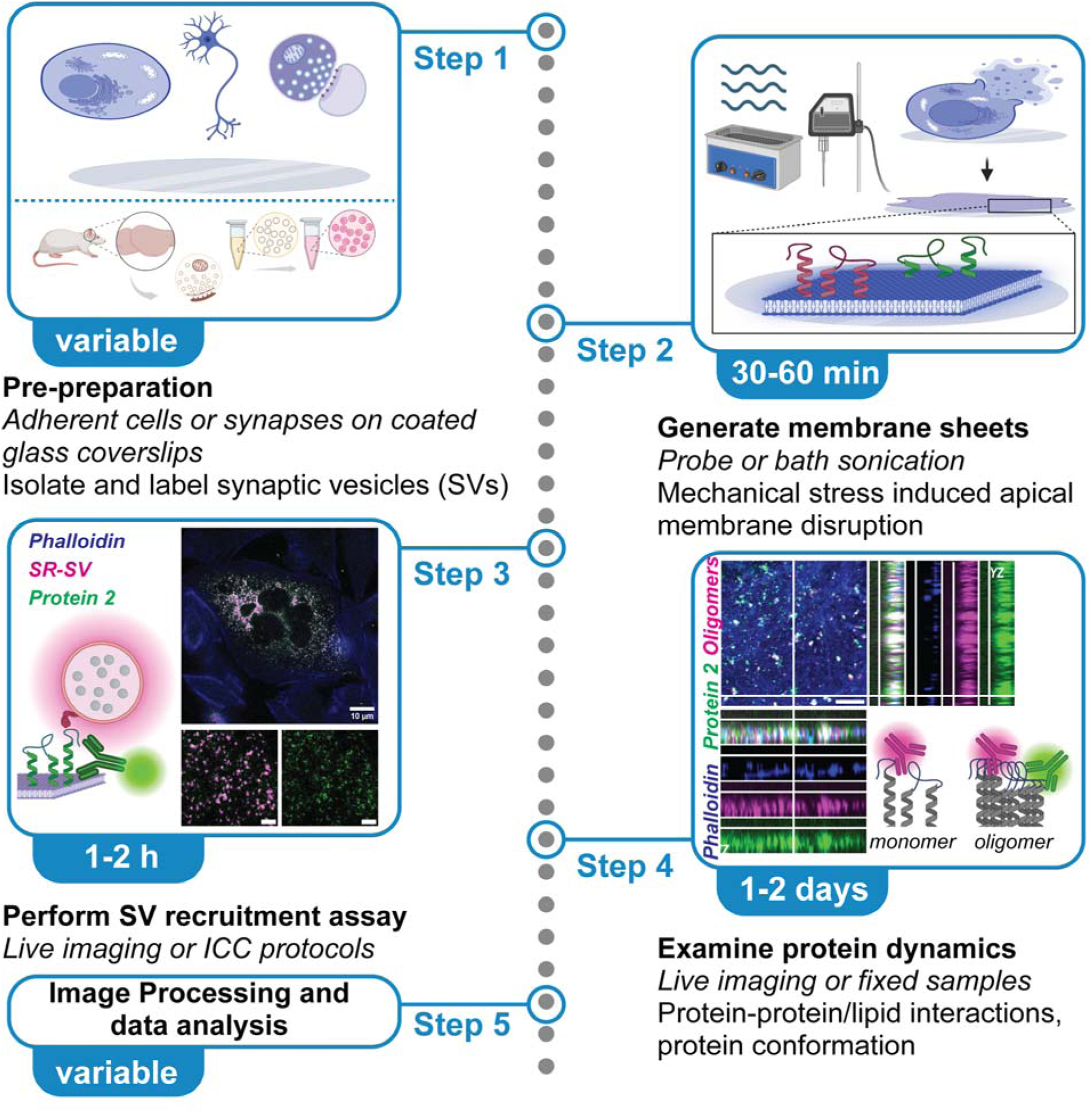

## Before you begin

This protocol describes an adaptation of the previously published single-molecule pull-down (SIM-Pull) assay for imaging synaptic protein complexes and membrane-associated synaptic vesicle recruitment using total internal reflection fluorescence (TIRF) microscope.^1^ While the previous protocol employed antibody-functionalized glass coverslips for single-molecule capture, the present protocol utilizes sonication-derived plasma membrane sheets to preserve the native organization of membrane proteins and lipids while exposing the cytosolic face of the plasma membrane.

The TIRF microscope configuration, imaging hardware, sample imaging chambers, and syringe pump-driven perfusion system have been described in detail previously.^1^ Here, the protocol is described using cells cultured on 25 mm glass coverslips mounted in a manual imaging chamber. However, the workflow can be readily adapted to alternative imaging chambers, microscope platforms, or perfusion systems with appropriate optimization.

### Innovation

Plasma membrane sheets provide a versatile platform for studying membrane-associated processes in a near-native environment.^2–4^ Membrane sheets derived by mechanical disruption (sonication) preserve the organization of endogenous membrane proteins and lipids while exposing the cytosolic face of the plasma membrane. Unlike reconstituted membrane systems^5,6^ or antibody-functionalized surfaces,^1,7–11^ this method preserves endogenous membrane lipid and protein composition; enabling quantitative analysis of membrane-associated interactions under near-physiological conditions.^2–4^

Here, we describe an optimized workflow for generating plasma membrane sheets from cultured cell lines, primary neurons, and isolated synaptosomes. We demonstrate their application to live imaging of membrane proteins, analysis of protein-protein interactions, and quantitative membrane-associated synaptic vesicle recruitment. The protocol is readily adaptable to virtually any adherent cell type, allowing researchers to transiently or stably express recombinant proteins, introduce disease-associated variants/mutations, or study endogenous proteins within their native membrane environment prior to membrane sheet preparation. This enables direct investigation of protein localization, membrane organization, self-assembly, conformational changes, and membrane-associated protein-protein interactions involved in synaptic vesicle tethering and other cellular processes. Furthermore, by combining membrane sheets with fluorescently labelled native synaptic vesicles, the platform enables quantitative analysis of membrane-associated vesicle recruitment and tethering.

Adaptation of this protocol can further enable studies to investigate nanoscale organization or membrane proteins or protein-dependent remodeling of membrane architecture or lipid organization. The modular nature of this protocol makes it broadly applicable for studying diverse membrane proteins and cytosolic membrane-associated processes while preserving their near-physiological membrane context.

### Institutional permissions

Animals were handled and maintained according to the guidelines laid down by the Animal Welfare Body (AWB) (Instantie voor Dierenwelzijn IvD) in line with the animal experimentation policy within Radboud University and RadboudUMC; under the license/protocol numbers 2021-0040-001/002 to Dr. Anne-Sophie Hafner.

### Cell culture and transfection

#### Timing: [variable]

Culture and transfection of Chinese hamster ovary (CHO) cells, including the generation of transiently transfected and stable cell lines, have been described previously^12,13^

Culture of rat primary cortical neurons from embryonic day 18 (E18) embryos, and their maintenance, has been described previously.^12,13^

As membrane sheets are typically (but not necessarily) generated after protein expression, the choice of cell type, culture protocols and transfection method can be tailored to the researcher’s biological question; and does not affect the downstream membrane sheet preparation and analysis. This protocol is readily adaptable to any adherent cell types and is compatible with a wide range of transfection and genetic manipulation approaches, including transient transfection, stable expression and/or viral transduction.

### Isolation and labeling of synaptic vesicles from adult rat brain

#### Timing: [1 day, variable]

Isolation, purification, fluorescent labelling using SynaptoRed, and quality control of native synaptic vesicles (SVs and SR-SV) from adult rat forebrains have been described in detail in our previously published STAR Protocol.^1^ We recommend following this protocol without modification to obtain highly purified, fluorescently labelled SVs suitable for membrane sheet-based recruitment assays described here.

## Key resources table

| REAGENT or RESOURCE | SOURCE | IDENTIFIER |
| --- | --- | --- |
| Antibodies |  |  |
| APP-Aβ mid region- specific (clone: 4G8) | Biologend | 800708 |
| 4G8 -Biotin conjugated | Biologend | 800704 |
| 4G8 -Alexa 488 conjugated | Biologend | 800714 |
| Oligomer A11 Polyclonal Antibody | ThermoFischer | AHB0052 |
| Anti-STX1A (rabbit polyclonal) IgG (Clone: 230470F5) | ProteinTech | 83159-6-RR |
| Anti-STX1A/1B (mouse monoclonal) (clone: 2D3B5) | ProteinTech | 66437-1-Ig |
| Goat Anti-Rabbit IgG H&L (Alexa Fluor® 488) | Abcam | ab150077 |
| Goat anti-rabbit IgG (H+L) Highly Cross-Adsorbed Secondary Antibody, Alexa Fluor™ 568 | Invitrogen | A11036 |
| Goat anti-rabbit IgG (H+L) Highly Cross-Adsorbed Secondary Antibody, Alexa Fluor™ 647 | Invitrogen | A21245 |
| Goat anti-mouse IgG (H+L) Highly Cross-Adsorbed Secondary Antibody, Alexa Fluor™ 647 | Invitrogen | A21246 |
| Goat Anti-Mouse IgG H&L (Alexa Fluor® 488) | Abcam | ab150113 |
| Goat Anti-Mouse IgG H&L (Alexa Fluor® 647) | Abcam | ab150115 |
| Streptavidin, Alexa Fluor™ 488 conjugate | Biologend | 405235 |
| Streptavidin, Alexa Fluor™ 647 conjugate | Biologend | 405237 |
| Bacterial and virus strains |  |  |
| -- |  |  |
| Biological samples |  |  |
| -- |  |  |
| Chemicals, peptides, and recombinant proteins |  |  |
| Buffer, PBS (10X, Dulbecco's) powder pack | VWR | A0965.9010 |
| EDTA | CarlRoth | 1E23.1 |
| Synaptic Protein Extraction Reagent (Syn-PER™) | ThermoFischer Scientific | 87793 |
| Protease Inhibitor Cocktail Set III, EDTA-Free - Calbiochem | Sigma Aldrich | 539134-1ML |
| PhosSTOP™ | Sigma-Roche | 4906845001 |
| HEPES | Sigma Aldrich | H4034 |
| Sucrose | Sigma Aldrich | S0389 |
| DMSO | Sigma Aldrich | 20-139 |
| Glycerol | VWR-Avantor | 7044.1000 |
| Tris-HCl | ThermoFischer Scientific | 228030010 |
| NaCl | Carl Roth | 3957.2 |
| NP-40 | bioPLUS | 22040045 |
| Sodium deoxycholate | Sigma Aldrich | D5670 |
| SDS (pellet form) | Carl Roth | CN30.3 |
| MgCl<sub>2</sub> | VWR Avantor | 7791-18-6 |
| KCl | VWR Avantor | 1952276-71-9 |
| Trypsin (cell culture grade) | Roche and Gibco | 03358658103 (R);<br>15400-054 (G) |
| Bovine Serum Albumin (BSA) Fraction V, NZ-Origin | Carlroth | 8076,4 |
| Gibco™ Horse Serum, New Zealand origin | ThermoFischer Scientific | 16050130 |
| Normal Goat Serum | Abcam | ab7481 |
| Basal Medium Eagle | Thermo Fisher | 21010046 |
| BrainPhys™ medium | STEMCELL | 05790 |
| NeuroCult™ SM1 | STEMCELL | 05711 |
| B27 Plus | Gibco | A3582801 |
| Penicillin-Streptomycin | Gibco | 15140122 |
| NEAA | Thermo Fisher | 11140050 |
| Sodium pyruvate | Thermo Fisher | 11360070 |
| Ham's F-12 | Gibco | 11765047 |
| Fetal bovine serum | Thermo Fisher | A5209501 |
| Poly-D-lysine | Sigma | A-003-E |
| Lipofectamine 2000 | Thermo Fisher | 11668027 |
| G418 | InvivoGen | ant-gn-1 |
| DAPT (GSI) | Cayman | #13197-5 |
| Mitotracker Green | MedChemExpress | HY-135056 |
| Lysotracker Red | Thermo Fisher | L12492 |
| SynaptoRed C2 (lipophilic dye, equivalent to FM®4-64) | Biotium | 70027 |
| WGA-Alexa 568 (glycosylated membrane protein marker) | Thermo Fisher | W56133 |
| Phalloidin-647 (filamentous actin marker) | Thermo Fisher | A22287 |
| Hoechst (Nuclei marker) | Thermo Fisher | 62249 |
| Critical commercial assays |  |  |
| BCA protein assay kit | ThermoFisher Scientific | 23227 |
| Deposited data |  |  |
| -- |  |  |
| Experimental models: Cell lines |  |  |
| CHO cells | ATCC | CCL-61 |
| Primary cortical neurons | Janvier (E18 rat embryos) | Sprague Dawley rat strain<br>(RjHan:SD//Janvier) |
| Experimental models: Organisms/strains |  |  |
| Rat (Females, 6-8-month-old) | Janvier | Sprague Dawley rat strain<br>(RjHan:SD//Janvier) |
| Oligonucleotides |  |  |
| -- |  |  |
| Recombinant DNA |  |  |
| Human APP695 (APP-FL) [Protein 1] | Addgene | 30137 |
| Human APP-CTFβ (C99) [Protein2] | Addgene | 30146 |
| Human C99<sup>ΔCT</sup> [Mutant protein] | Addgene | 30143 |
| Software and algorithms |  |  |
| Image Studio/ Image Studio Lite | LiCOR Bio | https://www.licorbio.com/image-studio |
| FIJI (ImageJ) | NIH | https://fiji.sc, RRID: SCR_002285 |
| GraphPad Prism 10 | PRISM Academy | https://www.graphpad.com, RRID: SCR_002798 |
| Other |  |  |
| Ultracentrifuge (for synaptosome isolation and SV pelleting) | Beckman Coulter Model Optima-90 (SN CXE14C01) Rotor SW41Ti and SW45Ti |  |
| Benchtop Ultracentrifuge (SV purification and labelling) | ThermoScientific X4R Pro |  |
| Probe sonicator | Sartorius Labsonic P 3mm probe |  |
| Bath sonicator | Branson B200 Ultrasonic Cleaner | 100-951-010 |
| Shaker | Linear: Laboshaker;<br>Orbital: Rotamax |  |
| Centrifuge | Eppendorf 5804R,<br>Heraeus-Fresco21 |  |
| High-precision glass coverslips (No. 1.5H, 25 mm) | VWR | 631-1584 |
| Thermomixer | Eppendorf | 5382000015 |
| Plate reader | Tecan | Model Spark M10 |
| Plasma cleaner (optional) | Diener Femto |  |
| Glass coverslips | VWR | 631-1577/79/84 |
| Quick release magnetic and non-magnetic imaging chambers | Warner Instruments | QR-40LP |
| Culture dish microincubator | Warner instruments | DH-40iL |
| Fusion 100-X Touch infusion syringe pump | KR Analytics |  |
| BD Plastipak Syringes Luer Lock | BD Syringes | 309630; 309628 |
| Fittings and tubing | IDEX/Darvin Microfluidics | CIL-P-642, CIL-P-857, CIL-P-678, BL-PTFE-1608-20, CIL-XP-245X, CIL-P-632, CIL-P-634, CIL-P-603, CIL-P-702, CIL-P-250X, CIL-P-259X |
| TIRF microscope | Nikon live-cell TIRF and epifluorescence microscope |  |

Antibody dilutions are provided as general working ranges optimized for this study. All primary and secondary antibodies should be used at dilutions optimized by the user for the respective application or according to the manufacturer’s recommendations.

## Materials and equipment

⍰ We recommend filtering the buffers using a 0.22 µm filter (indicated in the respective recipes). This would aid in getting rid of any small dust/particles etc. which could result in unwanted artefacts during imaging. Autoclaving water before use is not necessary.
⍰ It is recommended to use low-retention Eppendorf tubes to minimize protein/vesicle/dye loss.
⍰ Preparation of all additional buffers and solutions used throughout this protocol, including imaging, washing, blocking, and immunostaining buffers, has been described in detail in our previously published STAR Protocol^1^ and should be prepared accordingly unless otherwise specified.
⍰ TIRF microscope with specific attachments as described earlier for optimal resolution of single vesicle events. Laser power and exposure should be adjusted to minimize photobleaching while ensuring adequate signal-to-noise ratio (SNR).
⍰ FIJI (ImageJ) macros may be customized for automated vesicle detection and background subtraction.
⍰ Cytosolic buffer

• Cytosolic buffer
| Reagent | Final concentration (mM) | Amount – 1X (50 mL) |
| --- | --- | --- |
| HEPES | 25 |  |
| MgCl<sub>2</sub> | 1.5 |  |
| KCl | 10 |  |
| EDTA | 1 |  |
| **Total** | **n/a** | **50 mL** |

Adjust the pH to 7.4 using NaOH before bringing the solution to the final volume with distilled water. Store at 4°C for up to six months.
**Note:** Immediately before use, supplement the buffer with:
1 mM dithiothreitol (DTT)
1X protease inhibitor cocktail
You can also prepare a 10 mM DTT stock solution and store ready to use aliquots at -20°C. These aliquots could be stored indefinitely.

**CRITICAL:** Supplement the cytosolic buffer with 1 mM DTT and 1X protease inhibitor cocktail immediately before each experiment. The supplemented buffer should be prepared fresh for every membrane sheet preparation, kept ice-cold at all times, and used on the same day. Maintain all buffers, samples, and coverslips on ice throughout the procedure to minimize protein degradation and preserve membrane-associated protein complexes.

## Step-by-step method details

### Preparation of membrane sheets from adherent cells

#### Timing: [30-60 min]

This protocol describes the preparation of plasma membrane sheets from adherent cells cultured on 25 mm glass coverslips. The process can be performed either by the use of a probe sonication (brief sonication) or a bath sonicator (longer sonication) in ice-cold cytosolic buffer to achieve controlled mechanical disruption of the apical plasma membrane and preserve the adherent basal membrane attached to the glass surface. Both approaches rely on mechanical removal of the apical membrane, cytosol, nuclei, and intracellular organelles, while retaining basal plasma membrane sheets containing associated proteins and native lipid organization in a near-physiological context (Fig. 1A, 2A, 3A *schemes*).

**Figure 1.**
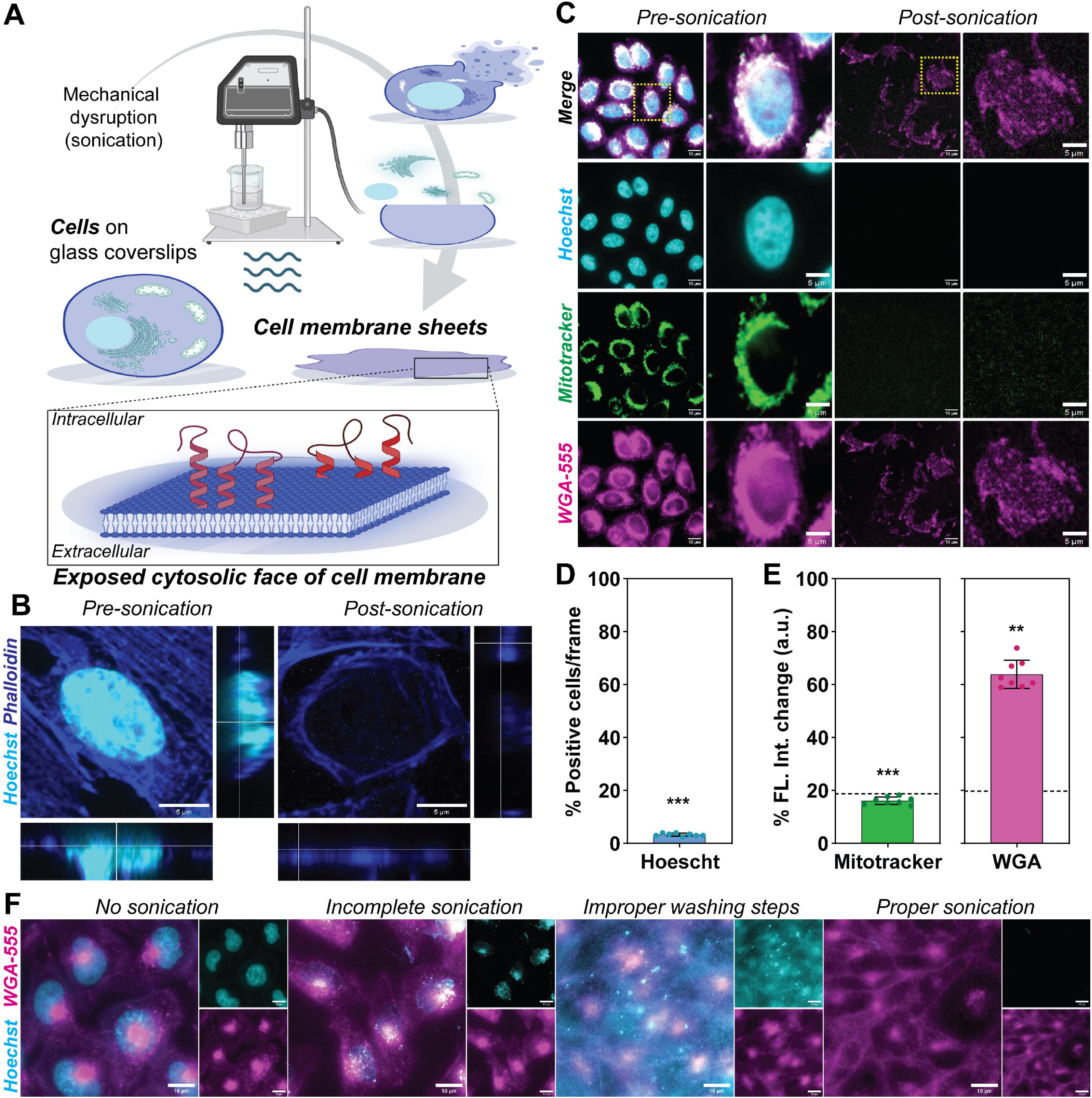
Preparation and validation of plasma membrane sheets generated by probe sonication. (A) Schematic overview of plasma membrane sheet preparation by probe sonication. Adherent CHO cells cultured on glass coverslips are immersed in ice-cold cytosolic buffer and subjected to brief probe sonication to mechanically remove the apical membrane, cytosolic contents, nuclei, and intracellular organelles while retaining the basal plasma membrane attached to the glass surface. This procedure exposes the cytosolic face of the plasma membrane, allowing direct access to the intracellular cytosolic domains of membrane-associated proteins while preserving their native membrane organization. (B) Representative fluorescence images of intact CHO cells (before sonication) and plasma membrane sheets (after probe sonication). Filamentous actin was labelled with phalloidin-647 (*blue*) and nuclei with Hoechst (*cyan*). Successful membrane sheet preparation results in retention of the basal plasma membrane while efficiently removing nuclei. Orthogonal views confirm the absence of nuclear material following sonication. Scale bar, 5 μm. (C) Representative fluorescence images of intact CHO cells before and after probe sonication. Nuclei were labelled with Hoechst (*cyan*), mitochondria with MitoTracker (*green*), and glycosylated plasma membrane proteins with WGA-555 (*magenta*). Following sonication, plasma membrane sheets remain attached to the coverslip while intracellular organelles are efficiently removed. Scale bars, 10 μm; zoomed-in images, 5 μm. (D) Quantification of the percentage of Hoechst-positive nuclei after membrane sheet preparation by probe sonication, normalized to values before sonication (100%). (E) Quantification of MitoTracker (*green*) and WGA-555 (*magenta*) fluorescence coverage before and after probe sonication, demonstrating efficient removal of intracellular components while preserving membrane-associated proteins. The dotted line indicates the background fluorescence level. Data are presented as mean ± SEM; N = 2. Statistical significance was determined using a two-tailed unpaired Student’s t-test. (F) Representative images depicting ideal and non-ideal images of membrane sheets post probe sonication. Incomplete sonication results in retention of nuclear material as well as intracellular organelles, wherein improper washing steps, or multiple usage of cytosolic buffer; resulting in accumulation of debris. Nucleus stained with Hoechst and glycosylated membrane proteins stained with WGA-555.

**Note:** The optimal sonication method and duration should be empirically determined for each cell type and experimental application.

1. Prepare cells and cytosolic buffer
  a. Culture the adherent cell type of interest on 25 mm glass coverslips until they reach approximately 70%-80% confluency.
  **Note:** Cells should be well attached and evenly distributed across the coverslip. Over-confluent cultures may generate overlapping or multilayered membrane sheets, whereas low-density cultures may yield insufficient membrane area for downstream analysis.
Prepare ice-cold cytosolic buffer and supplement it immediately before use with 1 mM DTT and 1X protease inhibitor cocktail. **CRITICAL:** The supplemented cytosolic buffer should be prepared fresh for every membrane sheet preparation kept ice-cold throughout the experiment and used on the same day.
2. Preparation of membrane sheets by probe sonication (Fig. 1A, *scheme*)
  a. Place 40-50 mL of supplemented ice-cold cytosolic buffer into a clean 50 mL glass beaker (Fig. 1A *scheme*).
  b. Transfer the coverslip containing adherent cells into the beaker with the cell-facing side oriented upward.
  c. Place the beaker on ice and allow the coverslip to equilibrate in cytosolic buffer for 1-2 min.
  d. Position the sonicator probe approximately 0.5-1 cm above the coverslip surface without touching the coverslip.
  **CRITICAL:** Adjust the volume of the buffer in the beaker, accordingly, ensuring that the coverslip is fully submerged in cytosolic buffer and does not touch the sonicator probe directly.

e. Sonicate the coverslip using using 35 mW output power (3 sonication pulses of 0.5 s each).

**CRITICAL:** Maintain the beaker on ice throughout sonication to prevent heating and excessive membrane disruption.

**Note:** Sonication intensity and cycle number may require optimization depending on the sonicator model, probe size/diameter, cell type, cell density, and coverslip coating as well as the protein of interest. Proper sonication procedure will result in membrane sheets devoid of nuclear and intracellular organelles as depicted in Fig. 1B, C and D-E. The decrease in the intensity of glycosylated membrane proteins observed in this case (Fig. 1E, *magenta*) could be due to loss of organelle membrane proteins as well as sonication induced mechanical stress. Under-sonication results in incomplete removal of cellular contents, whereas over-sonication leads to fragmentation or complete loss of membrane sheets (Fig. 1F).

f.Carefully remove the coverslip from the beaker using fine forceps and immediately transfer to a clean multi-6-well plate containing ice-cold cytosolic buffer.
g. Wash the coverslip gently 2-3 times with ice-cold cytosolic buffer to remove detached cellular debris.

**Note:** To minimize contamination from cellular debris and released intracellular components, process one coverslip at a time and do not use the same cytosolic buffer for more than three coverslips. Use freshly prepared ice-cold cytosolic buffer for each sample type. Replace the buffer with freshly supplemented cytosolic buffer after every third sample or sooner if visible debris accumulates. Improper washing or multiple usage of cytosolic buffer results in unclean membrane sheets unsuitable for single molecule imaging experiments (Fig. 1F).

**Figure 2.**
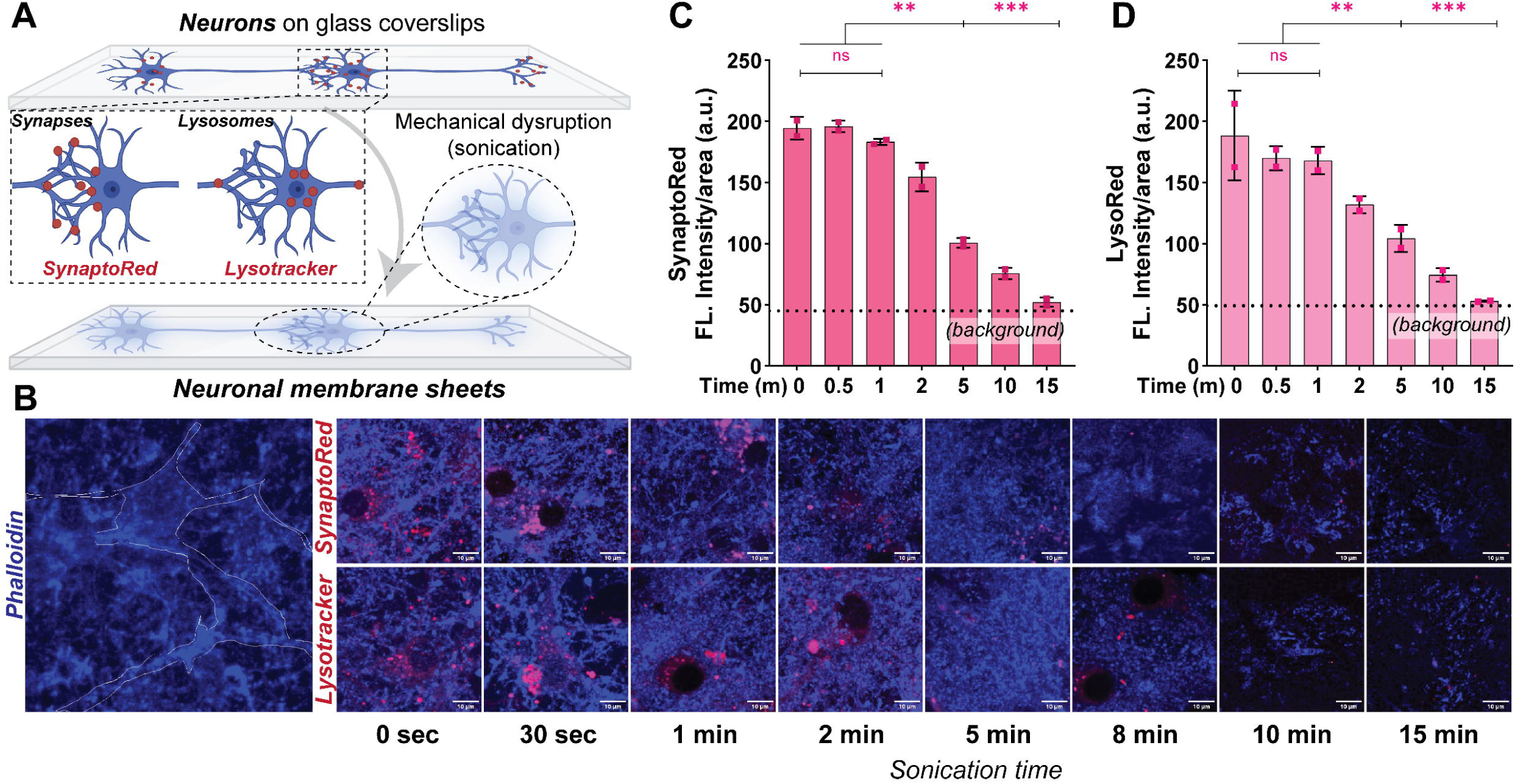
Optimization of membrane sheet preparation from primary neuronal cultures using bath sonication. (A) Schematic overview of membrane sheet generation from primary neuronal cultures. Mature DIV21 primary neurons cultured on poly-D-lysine (PDL)-coated glass coverslips were immersed in ice-cold cytosolic buffer and subjected to bath sonication for increasing time intervals to optimize membrane sheet preparation. (B) Representative fluorescence images of primary neuronal cultures before sonication (0 min) and following bath sonication for the indicated durations (0.5, 1, 2, 5, 10, and 15 min). Filamentous actin associated with the plasma membrane was labelled using phalloidin-647 (*blue*). Presynaptic terminals were labelled with SynaptoRed (*magenta, upper panels*), while lysosomes were labelled using LysoTracker Red (*magenta, lower panels*). Progressive bath sonication efficiently removes synaptic connections and intracellular organelles while preserving membrane sheets adhered to the glass coverslip. Scale bars, 10 μm. (C) Quantification of the percentage fluorescence coverage of SynaptoRed-positive synaptic puncta (**C**) and LysoTracker-positive lysosomes (**D**) following increasing bath sonication times, demonstrating progressive removal of synaptic structures and intracellular organelles. Data are presented as mean ± SEM; N = 2. Statistical significance was determined using one-way ANOVA with the appropriate post hoc multiple-comparison test (GraphPad Prism).

3. Bath sonication derived membrane sheets (Fig. 2A, 3A, *schemes*)

**Note:** Here, we used either isolated synaptosomes or mature primary cortical neurons (DIV21). Protocol for Synaptosome isolation and neuron preparation and maintenance has been described in detail elsewhere.^1,12,13^ Primary neuronal cultures should be maintained according to standard culture protocols until the desired developmental stage. Synaptosomes should be freshly isolated and allowed to adhere uniformly to gelatinated coverslips by centrifugation at 4000 x g for 15-30 min at 4°C before sonication. Details on gelatinization of coverslips have been described previously.^1,14^

a. Transfer each coverslip into a culture dish or glass container containing sufficient ice-cold cytosolic buffer to completely immerse the sample.
b. Place the container inside an ice-cold ultrasonic water bath. Fill the bath sonicator with ice pellets.

**Note:** To maintain an ice-cold bath sonicator, fill with 1:1 cold water and ice pellets.

**Tip:** Addition of NaCl lowers the freezing point and helps maintain the bath temperature close to 0°C throughout sonication.

c. Sonicate the samples for the desired duration to determine the optimal membrane sheet preparation conditions for the specific sample type as per user requirement and experimental application.

**Note:** Representative optimization of bath sonication conditions for primary neuronal cultures and isolated synaptosomes is shown in Fig. 2 and 3, respectively. Here, we have depicted data for increasing time intervals (e.g., 0.5, 1, 2, 5, 10, and 15 min, for primary neurons, Fig. 2B and 0.5, 1, 2, and 5 min, isolated synaptosomes, Fig. 3B). The optimal sonication time should preserve continuous membrane sheets and its protein components (Fig. 3D) while efficiently removing cytosolic components and intracellular organelles (Fig. 2C-D, Fig. 3C).

**CRITICAL:** Maintain the bath temperature between 0°C and 4°C throughout sonication by performing the procedure on ice or by regularly replacing the water with ice-cold water. Excessive heating will compromise membrane integrity and protein preservation.

d. Remove the coverslips using fine forceps from the container and gently wash 2-3 times with fresh ice-cold cytosolic buffer to remove any detached cellular material and debris.
e. Assemble the coverslips into the imaging chamber and maintain the membrane sheets in imaging buffer or cytosolic buffer until further processing.

Note: SynPull^15^ enables high-resolution characterization of protein aggregates within intact synaptosomes using single-molecule pull-down and super-resolution microscopy. In contrast, the membrane sheet platform described herewith enables functional investigation of specifically membrane protein organization, its interactions or conformational changes, and protein-mediated synaptic vesicle recruitment in a near-native/physiological membrane environment.

**Figure 3.**
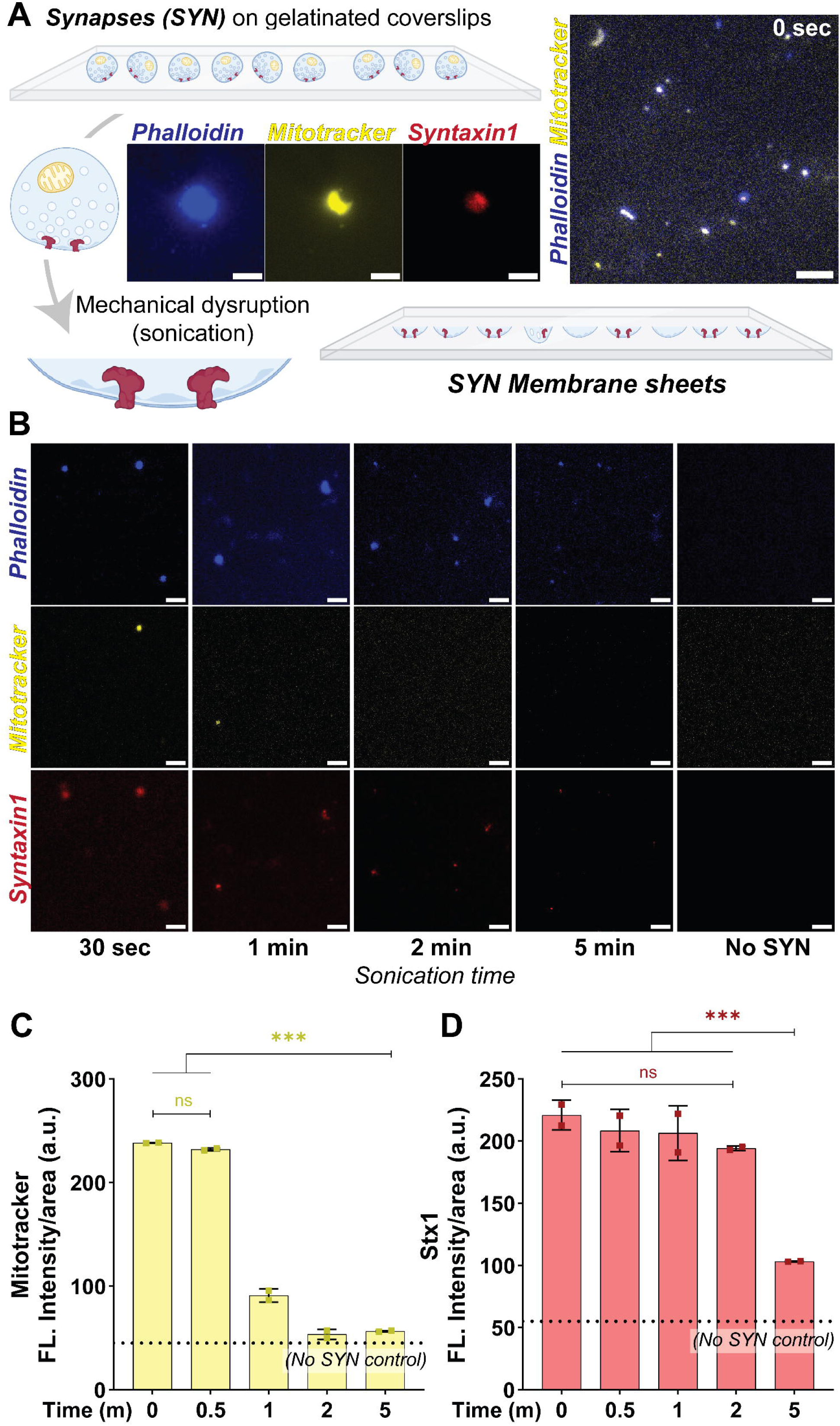
Optimization of membrane sheet preparation from isolated synaptosomes using bath sonication. (A) Schematic overview of membrane sheet generation from isolated synaptosomes. Freshly isolated synaptosomes adhered on gelatin-coated glass coverslips, were immersed in ice-cold cytosolic buffer, and subjected to bath sonication for increasing time intervals to optimize membrane sheet preparation. (B) Representative fluorescence images of adhered synaptosomes before sonication (0 min) and following bath sonication for the indicated durations (0.5, 1, 2, and 5 min). Synaptic membranes were visualized using phalloidin (*blue*), mitochondria were labelled with MitoTracker (*yellow*), and the presynaptic membrane protein Syntaxin-1 (STX1) was visualized by live immunolabelling (*red*). Together, these markers were used to evaluate preservation of synaptic membranes and the progressive removal of intra-synaptic organelles during sonication. Gelatin-coated coverslips without adhered synaptosomes (No SYN) were included as negative controls to assess nonspecific fluorescence arising from the gelatin coating. (C) Quantification of the percentage fluorescence coverage of MitoTracker-positive mitochondrial puncta (C) and Syntaxin-1-positive membrane puncta (D) following increasing bath sonication times. Bath sonication progressively removes intra-synaptic organelles while preserving membrane-associated synaptic proteins on the membrane sheets. Data are presented as mean ± SEM; N = 2. Statistical significance was determined using one-way ANOVA followed by the appropriate post hoc multiple-comparison test (GraphPad Prism).

4. Preparation of membrane sheets for live imaging and downstream applications

a. Assemble the coverslip into an imaging chamber with the membrane sheet-facing surface (cytosolic face) oriented upward.
b. Add imaging buffer or cytosolic buffer to completely immerse the membrane sheets.

**Note:** The imaging chamber used in this study accommodates a maximum imaging volume of 500 μL.

c. Proceed immediately to membrane sheet validation, immunostaining, or synaptic vesicle recruitment assays as described in the next steps.
d. During the initial optimization experiments, we recommend including non-sonicated control coverslips alongside sonicated samples to evaluate the efficiency of membrane sheet generation. Compare the two conditions by staining for nuclear (Hoechst) and intracellular organelle markers (Mitotracker, Lysotracker, SynaptoRed etc.) to confirm efficient removal of cellular contents while preserving intact plasma membrane sheets using phalloidin staining. Membrane integrity can additionally be assessed using Wheat Germ Agglutinin (WGA), which labels glycosylated plasma membrane proteins.
e. If the protein of interest is expressed as a fluorescent fusion protein (e.g., GFP, mCherry, HaloTag, SNAP-tag), membrane sheets can be directly imaged without further staining, making this approach particularly suitable for live imaging of protein localization and dynamics.
f. If the protein of interest is not fluorescently tagged, its localization and membrane retention can be examined by live immunolabelling using antibodies directed against an accessible epitope prior to membrane sheet preparation or by immunostaining following fixation, depending on the experimental objective.

### Live imaging of synaptic vesicle recruitment *via* protein-protein interactions on membrane sheets

#### Timing: [1-2 h]

This assay enables quantitative analysis of synaptic vesicle tethering mediated by endogenous or genetically expressed membrane-associated proteins while preserving their native membrane organization. This assay enables quantitative analysis of synaptic vesicle tethering mediated by endogenous or genetically expressed membrane-associated proteins while preserving their native membrane organization.^13^

**Note:** Preparation, purification, fluorescent labelling, quality control of synaptic vesicles, and the flow-through washing system have been described in detail previously.^1^ Only modifications specific to membrane sheet assays are described below.

5. Synaptic vesicle recruitment assays (Fig. 4A, *scheme*)

a. Place the imaging chamber on the microscope stage and identify suitable membrane sheet regions using the membrane marker, expressed fluorescent protein, or brightfield signal.
b. Acquire baseline images of good membrane sheet regions before addition of synaptic vesicles.
c. Add freshly prepared fluorescently labelled synaptic vesicles directly to the imaging chamber.
d. As described previously, SynaptoRed-labelled synaptic vesicles provided robust and reproducible fluorescent signals for live vesicle recruitment assays.

**Figure 4.**
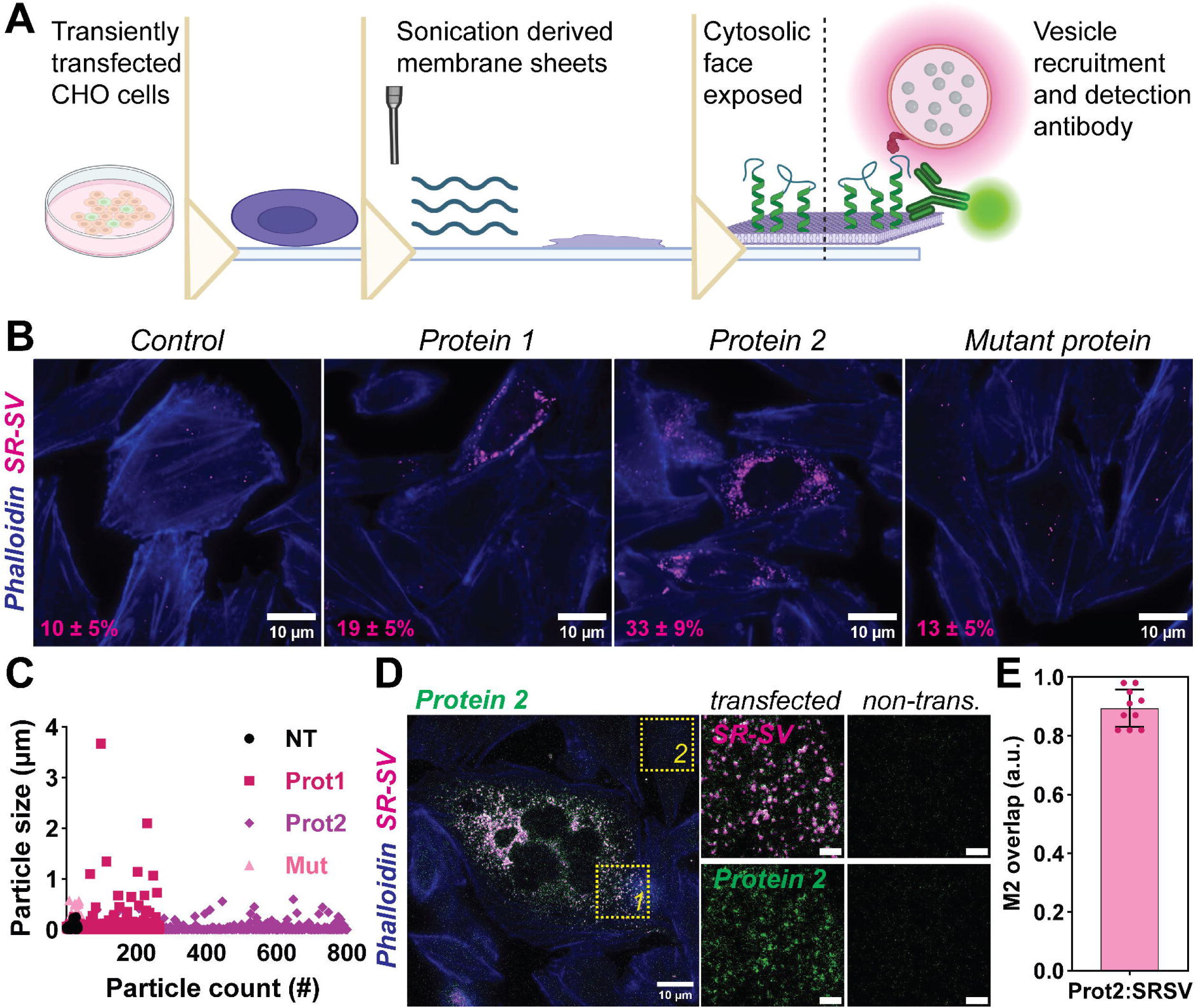
Live imaging of membrane-associated synaptic vesicle recruitment using plasma membrane sheets. (A) Schematic overview of the membrane sheet-based synaptic vesicle recruitment assay. Freshly prepared membrane sheets expressing the protein of interest were incubated with fluorescently labelled native synaptic vesicles (SynaptoRed-labelled synaptic vesicles; SR-SVs), followed by gentle washing using a syringe pump-driven flow-through system to remove unbound vesicles prior to fluorescence imaging. (B) Representative fluorescence images showing recruitment of SynaptoRed-labelled synaptic vesicles (SR-SVs) onto membrane sheets expressing the protein of interest. Membrane-associated synaptic vesicles appear as discrete fluorescent puncta associated with phalloidin-positive membrane sheet regions. Scale bars, 10 μm. (C) Representative particle analysis showing the distribution of membrane-associated SR-SV puncta according to particle size and particle number. This analysis can be used to compare synaptic vesicle recruitment between wild-type and mutant proteins, pharmacological treatments, antibody-blocked or protease-treated synaptic vesicles, or other experimental perturbations affecting membrane-associated protein interactions. (D) Representative workflow for quantitative image analysis. Protein-positive membrane regions were segmented to generate a mask for the protein of interest. Membrane-associated SR-SV puncta were identified using threshold-based particle analysis and quantified after normalization to either the total membrane sheet area or the protein-positive membrane area. Scale bars, 10 μm. (E) Colocalization analysis demonstrating that membrane-associated SR-SV puncta localize predominantly to protein-positive membrane regions. Colocalization was assessed using object-based analysis following particle segmentation. This illustrates the suitability of the assay for quantitative analysis of membrane-associated protein-mediated vesicle tethering. Images are representative of at least three independent experiments. Data are presented as mean ± SEM.

**Note:** SynaptoRed fluorescence gradually decreases during repeated washing steps owing to dilution of unbound dye and, over prolonged experiments, partial vesicle rupture. To maximize signal intensity, minimize the imaging chamber volume wherever possible. Alternatively, users may fluorescently label specific synaptic vesicle proteins or distinct vesicle populations using genetically encoded fluorescent proteins or fluorescent antibodies depending on the biological application.

**CRITICAL:** Use freshly thawed synaptic vesicle preparations whenever possible. Multiple freeze-thaw cycles or prolonged incubation at room temperature reduce vesicle integrity and may decrease specific vesicle recruitment while increasing nonspecific background as also described earlier.

d. Incubate the membrane sheets with fluorescent synaptic vesicles for 20 min.
e. Wash the chamber gently using the syringe pump-driven flow-through system described previously^1^ to remove unbound synaptic vesicles while minimizing disturbance of membrane-bound vesicles.

**CRITICAL:** Membrane sheets should remain hydrated throughout the experiment. Avoid drying at any stage, as this will irreversibly damage/disrupt the membrane and increase nonspecific vesicle binding.

f. Acquire images of membrane-bound fluorescent synaptic vesicles using TIRF or other high-resolution fluorescence microscopy modalities.

**Note:** For live recruitment experiments, time-lapse imaging before, during, and after synaptic vesicle addition is recommended during assay optimization to verify that vesicle recruitment occurs progressively and is mediated by specific protein-protein interactions rather than non-specific membrane adhesion.

g. Quantify membrane-bound synaptic vesicles as fluorescent puncta normalized to membrane sheet area or protein-positive membrane area, as described in the Image analysis and quantification section.

**Note:** When transiently transfected cells are used, neighboring non-transfected cells within the same field of view can serve as internal controls for nonspecific vesicle binding. Vesicle recruitment can therefore be normalized to the average background recruitment measured on adjacent non-transfected membrane sheets, reducing variability between experiments.

h. This assay can be readily adapted to investigate diverse membrane-associated protein interactions by expressing recombinant proteins, disease-associated variants, or fluorescent biosensors, or by immunolabelling endogenous proteins on membrane sheets. Synaptic vesicle recruitment then serves as a functional readout of membrane-associated protein interactions. Immunocytochemistry procedures have been described previously.^1^

### Live imaging of membrane protein organization and conformational dynamics using membrane sheets

#### Timing: [1-2 h, variable]

This protocol describes live imaging of membrane-associated proteins on freshly prepared membrane sheets. Freshly prepared membrane sheets provide direct optical access to the cytosolic face of the plasma membrane while preserving the native organization of membrane proteins and lipids. This enables live imaging of protein localization, nanoscale organization, self-assembly, conformational dynamics, and membrane-associated protein interactions under near-native conditions.

6. Live imaging of protein dynamics in membrane sheets

a. Mount the imaging chamber onto the microscope stage and identify intact membrane sheet regions using transmitted light or membrane-specific fluorescent markers.

**CRITICAL:** Select membrane sheet regions that are intact, continuous, and free of residual nuclei or intracellular debris. Regions containing membrane folds, fragmented sheets, or detached cellular material should be excluded from quantitative imaging.

b. Visualize membrane proteins by live immunolabelling using fluorophore-conjugated antibodies directed against extracellular epitopes (see Immunocytochemistry protocol^1^)
c. Optionally, if the protein of interest is expressed as a fluorescent fusion protein or fluorescent biosensor, live imaging can be performed directly without antibody labelling. This approach is preferred for monitoring protein dynamics in real time because it avoids antibody-induced perturbations and additional washing steps.

**Note:** This platform is compatible with fluorescent protein fusions, conformation-sensitive antibodies, FRET- and BiFC-based biosensors, polarity-sensitive membrane dyes, fluorescent ligands, and live-cell compatible antibodies.

d. Acquire baseline images before initiating time-lapse acquisition, applying experimental treatments, or adding compounds of interest (e.g., ligands, peptides, proteins, ions, pharmacological agents, or imaging buffers).
e. Acquire time-lapse images using the desired acquisition interval and duration according to the biological process being investigated.

**CRITICAL:** Membrane sheets should remain fully immersed in imaging buffer throughout live imaging. Minimize laser power and exposure time to reduce photobleaching and phototoxic effects. Freshly prepared membrane sheets provide the best preservation of native membrane architecture and are therefore strongly recommended for live imaging experiments.

f. Quantify changes in protein localization, clustering, oligomerization, fluorescence intensity, molecular mobility, conformational state, or membrane distribution using image analysis methods appropriate for the experimental design.

**Note:** Depending on the experimental application, membrane sheets can be used to study protein trafficking, lateral membrane diffusion, self-assembly, oligomerization, protein-protein interactions, conformational changes, membrane domain organization, lipid remodeling, and dynamic responses to pharmacological or physiological stimulation while maintaining proteins within their native membrane environment.

**Figure 5.**
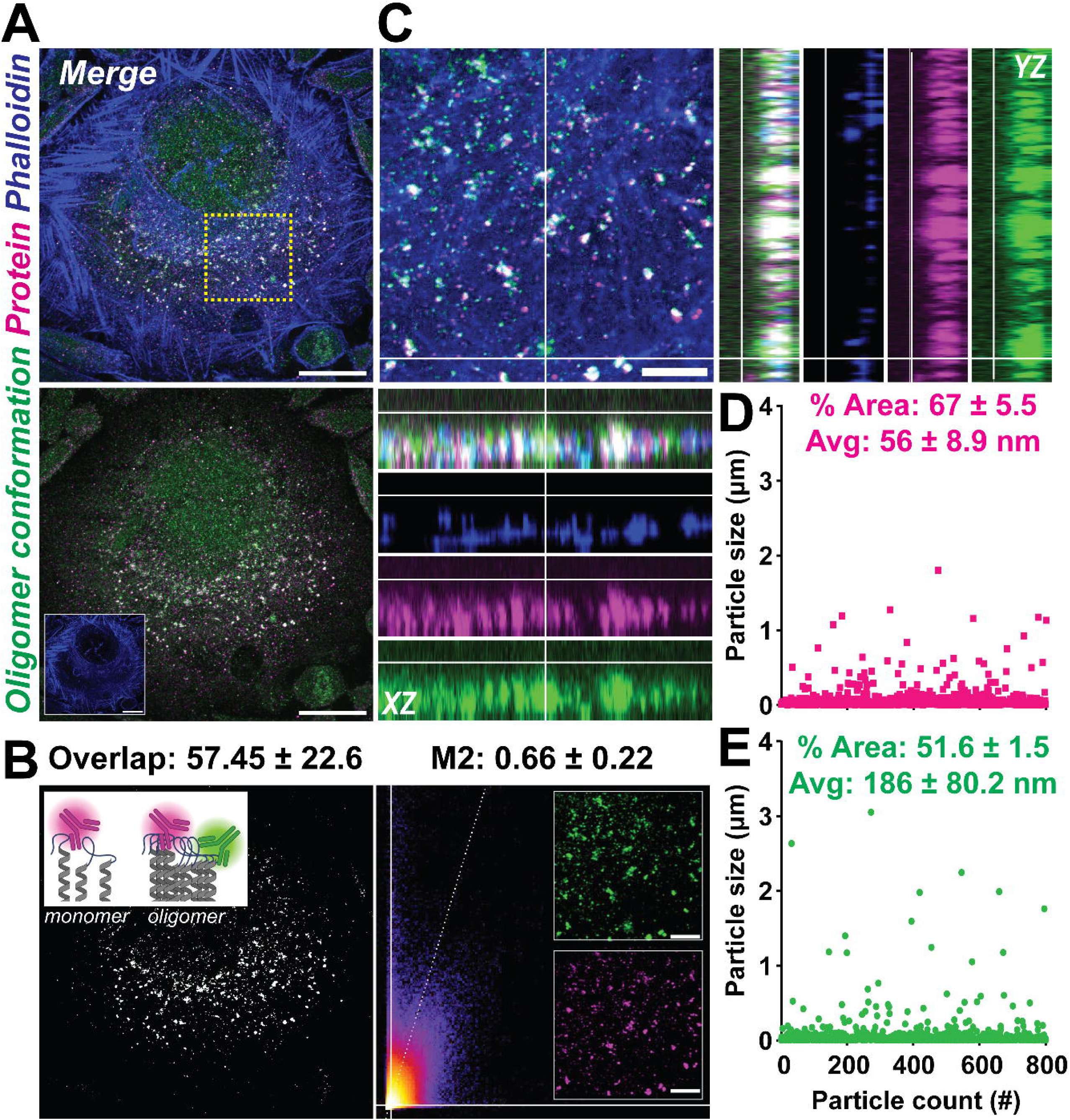
Immunofluorescence-based analysis of membrane protein organization and conformation on membrane sheets. (A) Freshly prepared membrane sheets were chemically fixed, blocked, and immunostained using antibodies recognizing either the total protein population or a conformation-specific epitope prior to fluorescence imaging. Representative fluorescence images of membrane sheets (phalloidin positive, *blue*) immunostained with antibodies recognizing the total protein of interest (*magenta*) and a conformation-specific epitope (*green*). Membrane sheets preserve membrane-associated proteins following fixation, enabling visualization of protein localization and conformational state while maintaining their native membrane organization. Scale bars, 10 μm. (B) Grayscale images illustrating the degree of colocalization between the total protein and conformation-specific antibody signals. Colocalization was assessed using the Manders M2 overlap coefficient, with representative overlap values and the corresponding intensity scatter plot shown alongside. Scale bars, 2 μm. (C) Representative higher-magnification images and orthogonal views illustrating the spatial distribution of the total protein and conformation-specific signals on membrane sheets. Orthogonal reconstructions demonstrate that both signals remain associated with the plasma membrane following membrane sheet preparation. Scale bars, 2 μm. (D) Quantitative analysis of the total protein (D) and conformation-specific (E) signals. Scatter plots depict the size distribution of individual fluorescent puncta, while accompanying bar graphs show the percentage fluorescence coverage and mean puncta area measured from membrane sheets following immunostaining. Images and data are representative of at least three independent experiments.

### Imaging membrane proteins on fixed membrane sheets

#### Timing: [2 days, variable]

Prepared membrane sheets may be chemically fixed when live imaging of membrane protein dynamics is not required. Fixed membrane sheets are suitable for immunofluorescence-based analysis of protein localization, protein-protein interactions, protein conformation, and membrane-associated protein organization.

7. Fixing membrane sheets

a. Prepare membrane sheets as described above and keep the coverslips in ice-cold cytosolic buffer.
b. Replace the cytosolic buffer with freshly prepared ice-cold fixation solution containing 4% paraformaldehyde (PFA) and 4% sucrose in PBS (fixing solution).
c. Incubate the membrane sheets in fixation solution for 10 min on ice.
d. Remove the fixing solution and wash the coverslips 4 times with ice-cold PBS.

Stopping point: You can store the samples for about 1 week at 4°C.

8. Immunostaining membrane sheets

a. Block the fixed membrane sheets using blocking buffer for 30-60 min at room temperature.

**Note:** Figure 5 illustrates a representative application of this protocol using cells expressing a membrane-associated protein that undergoes self-assembly into oligomers. Protein conformation was assessed using the oligomer-specific antibody A11 together with an antibody recognizing total protein.

**CRITICAL:** The use of permeabilizing agents such as TRITON X-100 should be avoided. If permeabilization is required, optimize the permeabilization conditions according to the detergent used and the protein of interest, as excessive permeabilization may disrupt membrane sheets or extract membrane-associated proteins.

b. Incubate the membrane sheets with primary antibodies diluted in blocking buffer according to user-optimized conditions or the manufacturer’s recommendations and incubate overnight at 4°C.
  i. Wash the coverslips 4 times with ice-cold PBS.
d. The washing steps could be performed on a laboratory shaker, with minimal/low shaking speed.
e. Incubate with species-appropriate fluorescent secondary antibodies diluted in blocking buffer for 1 h at room temperature, protected from light.
f. Wash the coverslips 4 times with ice-cold PBS.
g. Counterstain with suitable membrane markers (e.g., BODIPY dyes), cytoskeletal (phalloidin), or nuclear markers (DAPI or Hoechst), if required.
Coverslips may be imaged immediately in imaging buffer or mounted onto microscope slides using an appropriate antifade mounting medium for long-term storage prior to fluorescence microscopy.

**Note:** Fixed membrane sheets are particularly suitable for studying the retention, spatial organization, conformational state, and interaction-dependent clustering of endogenous or recombinant membrane proteins. Representative examples demonstrating analysis of total protein localization and conformation-specific antibody staining are shown in Figure 4.

**CRITICAL:** Avoid allowing membrane sheets to dry at any stage following fixation, as drying can increase nonspecific antibody binding and disrupt membrane morphology.

### Image analyses and quantification

The image analysis strategy should be adapted to the biological question. However, all quantifications within an experiment should be performed using identical image acquisition settings, thresholding parameters, and particle detection criteria.

#### Timing: [variable]

9. Quantification of synaptic vesicle recruitment (Fig. 4C)

A detailed workflow for quantifying synaptic vesicle recruitment events and assessing capture efficiency and protein-protein interaction specificity has been described previously.^1^ Only modifications specific to membrane sheet assays are described below.

a. Analyze acquired images using ImageJ/Fiji or custom scripts (e.g., MATLAB, Python), depending on user preference and experimental requirements.

**Note:** The workflow described below was developed using Fiji/ImageJ. Quantification of synaptic vesicle puncta using the Analyze Particles function has been described in detail previously.^1^

b. Split the image into individual channels corresponding to the membrane marker, protein of interest, and recruited synaptic vesicles.
c. Generate a membrane sheet mask using the phalloidin-positive, WGA-positive, or any membrane-selective fluorescent dye.
d. Threshold the membrane marker channel to define the total membrane sheet area.
e. Generate a second mask using the protein-of-interest channel, if recruitment is to be quantified specifically within protein-positive membrane regions.

**Note:** In our experiments, membrane sheets generated from transiently transfected cells were analyzed by generating a second mask corresponding to Protein 1- or Protein 2-positive membrane regions.

f. Threshold the synaptic vesicle channel and identify membrane-associated vesicle puncta using Analyze Particles.
g. Exclude particles larger than 0.5 μm to exclude large aggregates or nonspecific fluorescent debris.
h. Quantify vesicle recruitment as the number of synaptic vesicle puncta normalized either to the total membrane sheet area or specifically to the protein-positive membrane area (Fig. 4D), depending on the experimental design.
i. For transient transfection experiments, calculate vesicle recruitment using:
Recruitment (%) = (protein-positive membrane-associated SV puncta / total membrane sheet area) × 100
j. Export all raw measurements, including membrane area, protein-positive area, vesicle puncta number, puncta size, and fluorescence intensity.
k. Perform all quantifications blinded to experimental conditions whenever possible.

10. Analysis of protein conformation on membrane sheets (Fig. 5)

The membrane sheet platform can also be used to quantify membrane protein localization, organization, and conformational state using fluorescence intensity- and colocalization-based image analysis. Protein conformation can be assessed by dual immunolabelling using one antibody recognizing total protein and a second antibody specific for a conformational epitope, post-translational modification, or oligomeric state of the same protein.

a. Acquire fluorescence images of both the total protein channel and the conformation-specific antibody channel using identical imaging settings across all experimental conditions.

**CRITICAL:** Both antibodies should recognize non-overlapping epitopes of the target protein and be validated for simultaneous immunostaining. Imaging parameters, threshold values, and analysis settings should remain identical across all experimental conditions.

b. Generate a mask of the membrane sheet using a membrane marker (e.g., WGA or phalloidin) and restrict all subsequent analyses to this region of interest.
c. Threshold the total protein channel to identify membrane-associated protein-positive regions or puncta (Fig. 5A, B).
d. Measure the fluorescence intensity, puncta number, area, or integrated density of the total protein signal (Fig. 5D, E).
e. Apply identical thresholding parameters to the conformation-specific antibody channel and quantify the corresponding fluorescence signal.
f. Determine the relative abundance of the conformational protein species can be determined by normalizing the conformation-specific signal to the total protein signal on the same membrane sheet (Fig. 5D, E).
g. Optionally, perform colocalization analysis between the total protein and conformation-specific channels using Pearson’s correlation coefficient, Manders’ overlap coefficient, or object-based colocalization analysis (Fig. 5B).
h. Export fluorescence intensity, puncta number, puncta size, and colocalization measurements for statistical analysis.

**Note:** This analysis enables quantitative assessment of changes in protein conformation, oligomerization, or epitope accessibility while simultaneously accounting for differences in total protein expression. Representative examples of total protein and conformation-specific immunolabelling on membrane sheets are shown in Figure 5.

11. Data Interpretation and Statistical Testing

Note: Data should be collected from multiple membrane sheets across at least three independent biological replicates using identical image acquisition and analysis parameters.

a. Compare membrane-associated synaptic vesicle recruitment between untreated and control conditions to determine the specificity of membrane-associated vesicle tethering (Fig. 4C).
  i. Untreated versus antibody-blocked synaptic vesicles.

Quantify the reduction in recruited synaptic vesicles following pre-incubation of synaptic vesicles with function-blocking antibodies directed against synaptic vesicle surface proteins of interest.^1^

ii. Untreated versus trypsin-digested synaptic vesicles.

Compare vesicle recruitment before and after limited proteolytic digestion of synaptic vesicles to determine dependence of vesicle recruitment on intact vesicle surface proteins.^1^

b. Customize the assay according to the biological question by comparing membrane sheets expressing different proteins or protein variants.

Examples include

i. Wild-type versus disease-associated mutants (Fig. 5B)
ii. Wild-type versus deletion or domain mutants (Fig. 5B)
iii. Different protein isoforms (Fig. 5B)
iv. Pharmacological or genetic perturbations affecting membrane-associated proteins
v. Different cell types or membrane preparations

c. Statistical analysis should be carried out using GraphPad Prism or equivalent software. Data should first be tested for normality before selecting the appropriate statistical test. Comparisons between two groups may be analyzed using an unpaired or paired two-tailed Student’s *t*-test, whereas comparisons involving multiple groups should be analyzed using one-way or two-way ANOVA followed by an appropriate multiple-comparison post hoc test. Statistical significance should be defined according to the criteria established by the investigator and reported in the corresponding figure legends.

**Note:** To minimize experimental bias, membrane sheet preparation, image acquisition, thresholding, and particle analysis should be performed using identical parameters across all experimental groups. Whenever possible, image acquisition and quantification should be performed blinded to the experimental condition.

## Expected outcomes

Successful preparation of membrane sheets should result in continuous plasma membrane sheets firmly attached to the glass coverslip while efficiently removing the apical membrane, cytosolic contents, nuclei, and most intracellular organelles. Membrane sheets should exhibit strong membrane staining (e.g., WGA or membrane dyes) together with preserved cytoskeletal architecture, while nuclear and organelle markers should be markedly reduced or absent, indicating efficient mechanical disruption and removal of intracellular contents (Figs. 1-3).

Depending on the experimental application, membrane sheets can be used immediately for live imaging or chemically fixed for subsequent immunostaining (Fig. 5). Live membrane sheets enable visualization of membrane protein localization, lateral organization, clustering, self-assembly, conformational dynamics, and protein-protein interactions under near-native conditions. Fixed membrane sheets preserve membrane-associated proteins and are suitable for immunofluorescence-based analysis of protein distribution, conformation, and interaction-dependent organization (Fig.5).

When combined with fluorescently labelled synaptic vesicles, membrane sheets support robust and reproducible membrane-associated vesicle recruitment (Fig. 4). Specific vesicle tethering should predominantly occur on membrane sheets expressing the protein of interest and can be quantified as membrane-associated fluorescent puncta following removal of unbound vesicles. Appropriate negative controls, including non-transfected cells, antibody-blocked synaptic vesicles, or protease-treated synaptic vesicles, should display markedly reduced vesicle recruitment, confirming that vesicle capture is primarily mediated by specific protein-protein interactions.

Overall, this protocol provides a versatile and broadly adaptable platform for investigating membrane protein organization, conformational changes, protein-protein interactions, membrane remodeling, and functional membrane-associated processes while preserving the native plasma membrane environment.

## Quantification and statistical analysis

⍰ Images acquired by TIRF or other high-resolution fluorescence microscopy modalities can be analyzed using Fiji/ImageJ or equivalent image analysis software. Before quantitative analysis, raw images should be corrected for background fluorescence by subtracting the average background signal obtained from blank membrane sheet preparations or control coverslips imaged under identical acquisition settings. Appropriate controls should include camera offset (dark images), coverslip autofluorescence, nonspecific fluorescence arising from imaging buffers, and membrane sheets processed in the absence of fluorescent probes or synaptic vesicles.
⍰ Background fluorescence should be determined from 6-10 independent images for each coverslip preparation and subtracted from all experimental images prior to analysis.

**CRITICAL:** Background fluorescence should remain low and consistent throughout the experiment. Large variations in background fluorescence between coverslips prepared under identical conditions may indicate increased nonspecific fluorescence, membrane damage, or technical variability and should be carefully evaluated before proceeding with quantitative analysis.

⍰ For synaptic vesicle recruitment assays, membrane sheets should first be segmented using a membrane marker (e.g., phalloidin, WGA, or another membrane-selective dye) to define the total membrane sheet area. When recruitment is analyzed specifically on transfected cells or membrane regions expressing the protein of interest, an additional mask should be generated from the corresponding fluorescence channel. Synaptic vesicle puncta can then be detected using threshold-based particle analysis (e.g., Analyze Particles) or spot-detection algorithms such as TrackMate. In this study, threshold-based particle counting was implemented as a Fiji/ImageJ macro to enable automated batch processing of all acquired images.
⍰ Only fields of view containing intact membrane sheets with homogeneous illumination and free of obvious membrane folds, cellular debris, or imaging artefacts should be included for quantitative analysis. Vesicle recruitment should be expressed as the number of membrane-associated fluorescent puncta normalized to either the total membrane sheet area or the protein-positive membrane area, depending on the experimental design.
⍰ Likewise, membrane protein localization, clustering, or conformational changes may be quantified using fluorescence intensity measurements, puncta analysis, colocalization, or conformation-specific signal normalized to total protein abundance.
⍰ All quantitative analysis should be performed from at least 10-15 fields of view per experimental condition across a minimum of three independent biological replicates. When transient transfection is used, neighboring non-transfected membrane sheets may serve as internal controls to normalize vesicle recruitment and reduce inter-experimental variability.
⍰ Data acquisition should be discontinued if substantial membrane fragmentation, vesicle aggregation, vesicle lysis, or marked increases in background fluorescence are observed, as these may compromise quantitative analysis.
⍰ Statistical analyses should be performed using GraphPad Prism or equivalent software. Data should first be tested for normality before selecting the appropriate statistical test. Comparisons between two groups may be analyzed using paired or unpaired two-tailed Student’s t-tests, whereas comparisons involving multiple groups should be analyzed using one-way or two-way ANOVA followed by appropriate post hoc multiple-comparison tests. Data should be reported as mean ± SEM (or SD where appropriate), with statistical significance and sample sizes indicated in the corresponding figure legends.
⍰ Statistical analyses should be performed using GraphPad Prism or equivalent software. Data should first be tested for normality before selecting the appropriate statistical test. Comparisons between two groups may be analyzed using paired or unpaired two-tailed Student’s t-tests, whereas comparisons involving multiple groups should be analyzed using one-way or two-way ANOVA followed by appropriate post hoc multiple-comparison tests. Data should be reported as mean ± SEM (or SD where appropriate), with statistical significance and sample sizes indicated in the corresponding figure legends.

## Limitations

Although this protocol preserves the native organization of plasma membrane proteins and lipids considerably better than reconstituted membrane systems, it remains an ex vivo preparation generated by mechanical disruption of intact cells. Consequently, cytosolic components, intracellular organelles, soluble binding partners, and ATP-dependent cellular processes are largely removed during membrane sheet preparation. As a result, membrane sheets are best suited for investigating membrane-associated protein organization, protein-protein interactions, conformational changes, and vesicle tethering events rather than active intracellular trafficking or signaling processes that require an intact cellular environment.

The quality of membrane sheets is highly dependent on the efficiency of mechanical disruption. Sonication conditions should therefore be optimized for each cell type, culture density, and sonication instrument, as insufficient sonication may leave residual cytoplasmic material attached to the membrane, whereas excessive sonication may fragment or detach membrane sheets and reduce protein retention. Likewise, some membrane-associated proteins, particularly those that are only weakly associated with the plasma membrane or require intact cytoskeletal interactions, may be partially lost during membrane sheet preparation.

For live imaging experiments, membrane sheets should be prepared immediately before imaging, as prolonged idle durations may gradually alter membrane integrity and protein organization. Similarly, fixation and immunostaining conditions should be optimized for individual proteins because certain epitopes or protein conformations may be sensitive to fixation or detergent-based permeabilization.

An additional consideration is that membrane sheets are generated from the ventral plasma membrane attached to the glass coverslip. Previous studies have reported that cell adhesion to glass can locally influence plasma membrane organization, including lipid composition, membrane tension, phosphoinositide distribution, and the formation or stability of membrane nanodomains (lipid rafts). Consequently, the membrane organization observed using this protocol primarily reflects the adherent membrane surface rather than the entire plasma membrane. While this provides a highly reproducible and experimentally accessible membrane preparation, users should consider this potential spatial bias when interpreting protein localization, membrane organization, lipid-dependent interactions, or comparing results with those obtained from intact cells.

Finally, quantitative analyses should be performed on multiple membrane sheets acquired from independent biological replicates and across multiple fields of view. Users should avoid selecting only visually optimal membrane sheets, as membrane size, morphology, and protein distribution may vary between preparations. Acquiring images from randomly selected membrane sheets over a broad imaging area minimizes sampling bias and provides a more representative assessment of membrane-associated protein organization and functional vesicle recruitment.

## Troubleshooting

### Problem 1

#### Poor membrane sheet generation or excessive membrane fragmentation

The efficiency of membrane sheet preparation is highly dependent on the sonication conditions. Insufficient sonication may leave intact cells, nuclei, or intracellular organelles attached to the coverslip, whereas excessive sonication may fragment or completely remove membrane sheets. Cell type, culture density, coverslip coating, and sonicator performance can all influence membrane sheet generation.

### Potential solution

- Optimize the sonication power and duration for each cell type and experimental preparation.
- Perform initial optimization experiments using increasing sonication times or power settings while monitoring membrane integrity using plasma membrane (e.g., WGA or membrane dyes), cytoskeletal (phalloidin), nuclear (Hoechst/DAPI), and organelle-specific markers.
- Maintain all buffers and samples at 0-4°C throughout the procedure and process one coverslip at a time using freshly supplemented cytosolic buffer.
- Make sure to perform enough washing steps before proceeding to the next stages of the experiment.

### Problem 2

#### High variability between fields of view or biased quantification

Membrane sheets can vary in size, morphology, protein expression level, and degree of mechanical disruption across the same coverslip. Selecting only large, bright, or visually optimal membrane sheets may introduce acquisition bias and overestimate protein clustering or synaptic vesicle recruitment.

### Potential solution

- Define inclusion and exclusion criteria before image acquisition.
- Exclude fields of view containing fragmented membrane sheets, residual nuclei, large debris, membrane folds, or uneven illumination.
- Acquire images from multiple randomly selected regions across each coverslip and include at least 10-15 fields of view per condition.
- Whenever possible, perform image acquisition and quantification blinded to the experimental condition using identical thresholding and particle analysis parameters across all groups.

### Problem 3

#### Weak protein signal or high background immunofluorescence

Poor antibody penetration, inappropriate antibody concentrations, membrane drying, excessive fixation or permeabilization, and nonspecific antibody binding can reduce signal quality or increase background fluorescence. Some membrane-associated proteins may also be partially lost during membrane sheet preparation.

### Potential solution

- Optimize antibody concentration and incubation times for the protein of interest.
- Avoid allowing membrane sheets to dry during fixation or staining procedures.
- Unless required, avoid detergent-based permeabilization, as this may disrupt membrane sheets and can extract membrane-associated proteins.
- Include appropriate negative controls and acquire images using identical imaging settings across all experimental conditions.

### Problem 4

#### Low or highly variable synaptic vesicle recruitment

Reduced vesicle recruitment may result from poor membrane sheet quality, low expression of the protein of interest, damaged or aged synaptic vesicle preparations, excessive washing, or variability between independent synaptic vesicle isolations.

### Potential solution

- Verify membrane sheet integrity before beginning recruitment assays and confirm expression or localization of the protein of interest.
- Use freshly prepared or freshly thawed synaptic vesicles and minimize repeated freeze-thaw cycles.
- Perform all recruitment experiments using identical incubation and washing conditions and normalize vesicle recruitment to membrane area or to neighboring non-transfected membrane sheets when transient transfection is used.
- Acquire data from multiple fields of view and independent biological replicates to account for experimental variability

### Problem 5

#### Progressive loss of SynaptoRed fluorescence during imaging

Prolonged incubation or imaging of membrane-bound synaptic vesicles may result in a gradual reduction of SynaptoRed fluorescence. We have occasionally observed diffusion of the SynaptoRed signal into the membrane sheet during extended imaging periods, which may reflect partial membrane fusion, lipid dye transfer, or reduced vesicle integrity over time.

### Potential solution

- To minimize fluorescence loss, perform image acquisition immediately following the final washing step and avoid prolonged incubation of membrane sheets with SynaptoRed-labelled synaptic vesicles.
- Use freshly prepared synaptic vesicles, minimize unnecessary exposure to excitation light, and acquire images as rapidly as possible while maintaining sufficient image quality.
- If extended time-lapse imaging is required, consider using fluorescent labels directed against synaptic vesicle proteins rather than membrane lipid dyes.

## Resource availability

⍰ ***Lead contact****: Dr. Anne-Sophie Hafner*
⍰ ***Technical contact****: Dr. Akshay Kapadia*
⍰ ***Materials availability***: This study did not generate new, or unique reagents. All reagents are commercially available.
⍰ ***Data and code availability***: All datasets generated during this study are made available upon request to the authors.

## Acknowledgments

⍰ European Research Council (ERC) under the European Union’s Horizon 2020 research and innovation program - ‘MemCode’, grant 101076961 (AH)
⍰ The authors thank Elena Cijffers for providing images depicted in Fig. 1C; and all our lab members for fruitful discussions and helpful feedback.
⍰ The authors acknowledge the support from General Instrumentation - Microscopy Core facility at Faculty of Science, Radboud University, Nijmegen, Netherlands; and thank Dr. Jelle Postma for his assistance with the TIRF microscopy set-up.
⍰ Graphical abstract/Figure schematics were created using Biorender.com.

## Author contributions

AK conceptualized the project, performed and analyzed experiments and wrote the original draft of the manuscript. ASH supervised AK, acquired funding, and reviewed and edited the manuscript.

